# Trantrace: a reliable, traceable, and collaborative solution to streamline the translation of emerging biomedical resources

**DOI:** 10.1101/2020.07.01.181677

**Authors:** Min Su, Zhaohong Wang, Mengyu Li, Weixing Ye, Peter Chen, Mo Huang, Zili He, Chaoran Xia, Jie Cui, Rui Liu, Congmao Wang

## Abstract

**Background:** Precision medicine is gaining popularity in routine health care, which heavily relies on the interpretation of emerging biomedical databases and professional guidelines. Translation of those biomedical materials, that is crucial to deliver research discovery to health interventions in non-native English-speaking countries, remains a large amount of work for biomedical practitioners.

**Results:** We presented an application called Trantrace to facilitate the routine of large-scale long-running translation. Especially for the case that many people joined multiple translation projects, it has a rigorous task division and cooperation process: assignment, translation, review, release, and iteration. Trantrace empowers users to build own projects with different permissions, version control all the translation operation without specific skills, and further improve the translation efficiency through real-time task management tools.

**Conclusions:** By providing a working platform for collaborative translation and enabling long-time version correction, the Trantrace contributes towards the creation of a multilingual biomedical knowledge commons, which would be a valuable aid for personalized therapies. The source code is freely accessible at https://github.com/sgi-drylab/trantrace.

## Background

In December 2018, Genomics England accomplished the sequencing 100,000 whole genomes from 70,000 patients and family members, which may serve as a clarion call to integrate genome sequencing into routine medical care [1]. The Cancer Genome Atlas (TCGA), which collected more than 11,000 cases across 33 tumor types, is creating lasting value to elucidate the complex and multi-dimensional problems of human cancers [2], and efforts of harmonized and standardized analysis, such as the Pan-Cancer Atlas, are tried to improve the ability to diagnose, treat and prevent cancers. The Human Microbiome Project (HMP) has been carried out to reveal contributions of the microbiome and its dynamics changes to human health and disease[3]. As more and more national and international genome programs gradually come to an end, it would usher in a new era of personalized medicine and early diagnosis. The rapidly increasing human genotype-phenotype databases collect flooded information about genetic variations and their phenotypic effect based on expert curation of published and unpublished data, which would provide reliable and comprehensive knowledge about the gene and related clinical or research topics [4]. Furthermore, the American College of Medical Genetics (ACMG) and the Association of Molecular Pathologists (AMP) had published a joint consensus to standardize the interpretation of sequence variants [5]. The Clinical Pharmacogenetics Implementation Consortium (CPIC) has published a series of evidence-based guidelines to individualize the prescribing of drugs [6]. Translation of these biomedical resources is crucial to tackle language barriers and bring the latest clinical and molecular findings or research subjects to everyone’s health care.

Traditional file-based working with Microsoft may have many pitfalls, including auto-formatting, not traceable and permission problems. A screen of supplementary Excel files in 2016 reported that one in five genetic papers in top scientific journals contains errors [7]. Though the accuracy of machine translation is greatly improved, it still can not replace human translation when it comes to scientific databases [8]. Translation management systems (TMS) are usually charged and designed for short-term commerce translation or software localization, which is not suitable for constantly updated resources. Here, we created a free application, Trantrace, to accelerate the translation speed, improve the translation quality, and facilitate the translation cooperation in large-scale long-running biomedical database translation.

## Implementation

### The graphic user interface

Trantrace provides a graphical user interface (GUI) (Fig. 1) and has almost no requirement for user skills. The sidebar is composed of five functional modules. Dashboard summarizes the user’s workload and completion by categories. Project management is for the translation projects you created and the entries to be assigned. Translation management is for the entries that you’re about to translate or have already translated. Review management is for the translations you’re about to review or have already reviewed. The released projects is for the projects that have released versions and issues that occurred in released translations. My task, the yellow button in the lower right corner, updates the total number of unfinished tasks every minute.

**Fig. 1.**
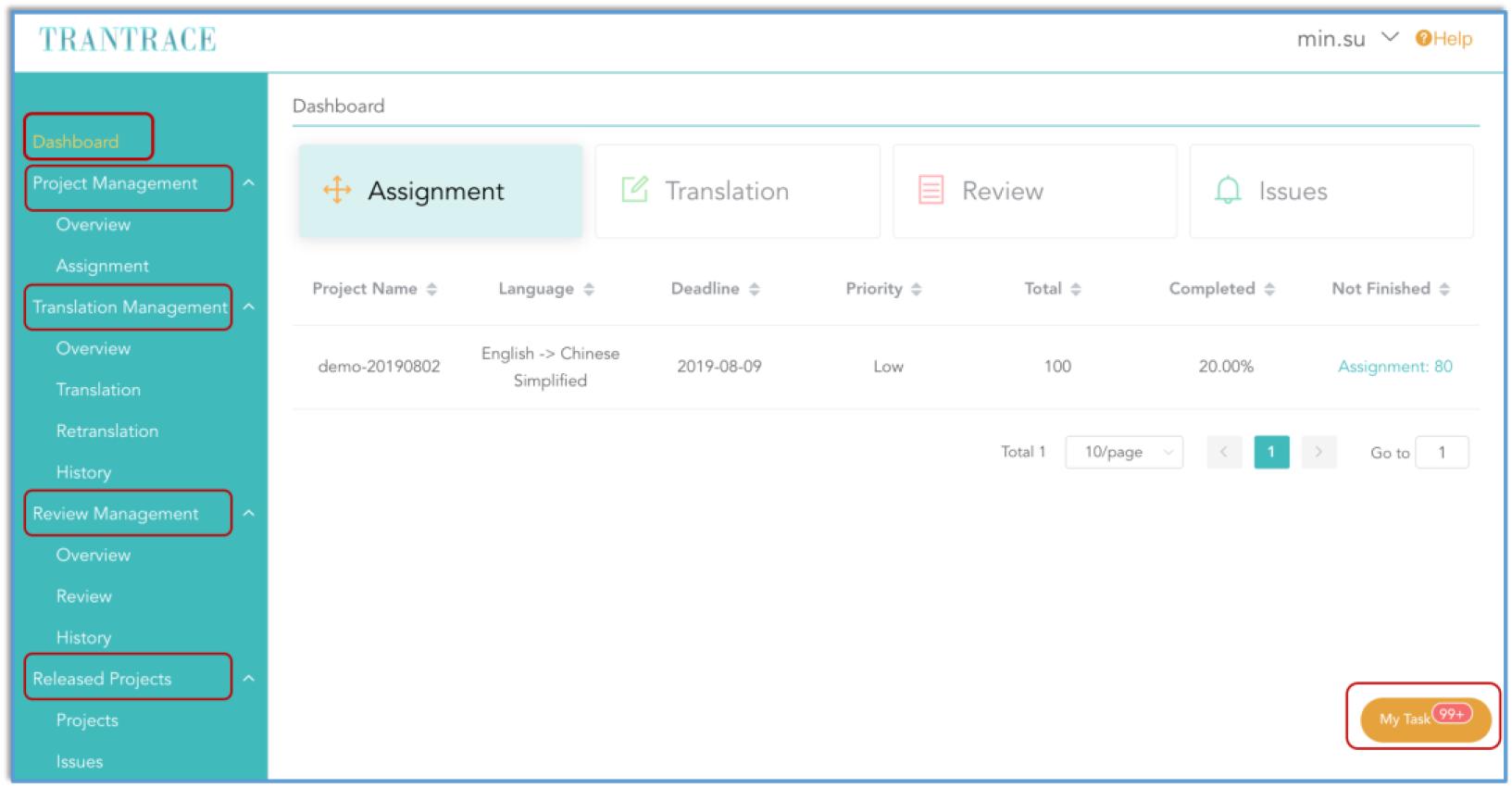
The GUI of Trantrace

### User permission

Permission in Trantrace depends on the project role instead of user identity. There are four different roles in one project as follows: owner, translator, reviewer, and guest (Fig. 2). The owner is in charge of the project and has the highest privilege to coordinate, including change setting, add member, file upload, task assignment, version release, and issue reply. The translator commits to translating assigned entries from the owner and re-translates failed entries from the reviewer. The reviewer, who decides whether to approve translations or not, is crucial for the translation quality. The guest has no obligation during the translation. All project roles can search released translations and open issues if found errors.

**Fig. 2.**
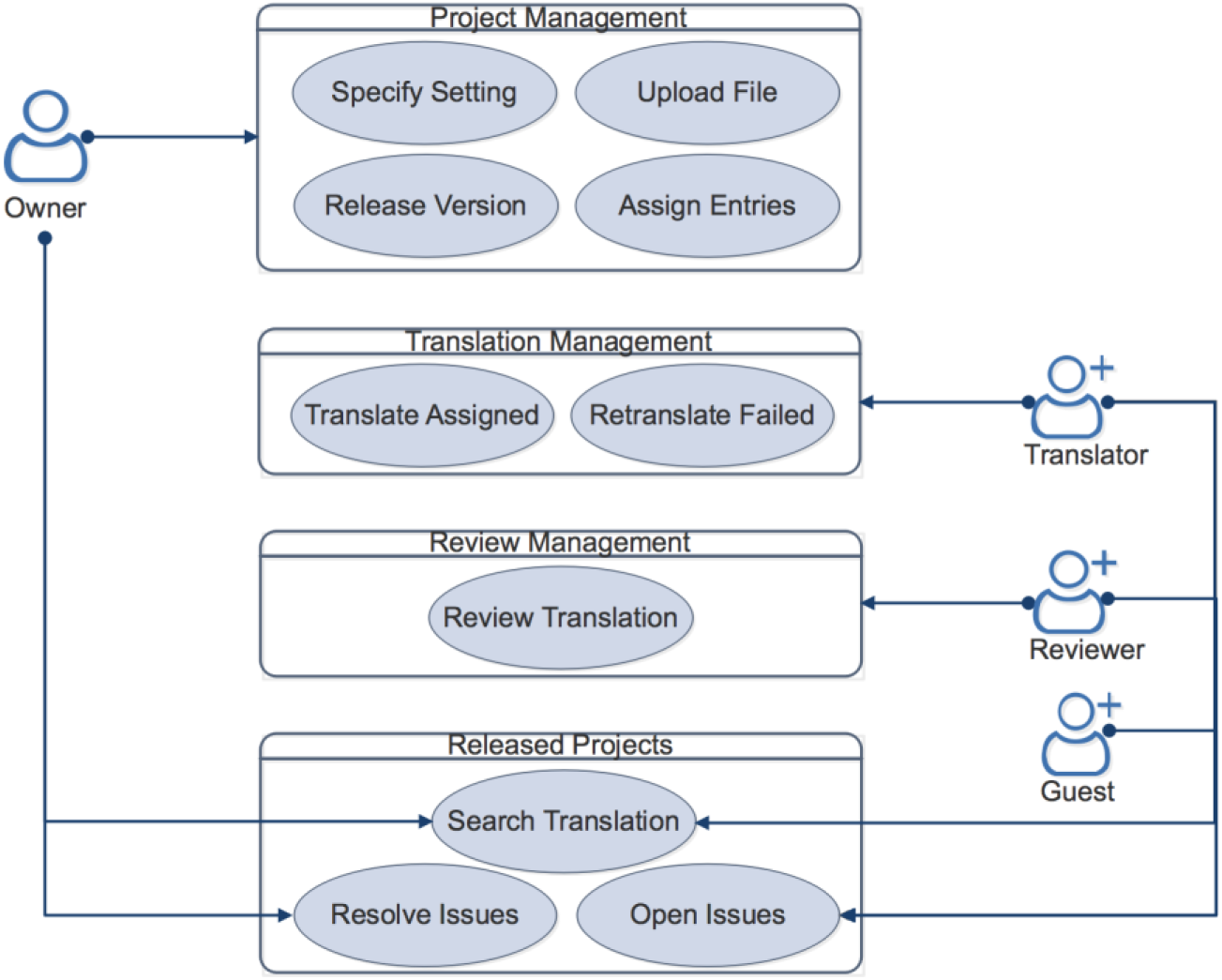
Project Role and Related Permission

### Data preparation

Raw data that download from websites is not suggested. Unrelated messages should be removed to reduce system load. The current support file format is a comma-delimited or tab-delimited text file and spreadsheet from Microsoft Excel. The first column, usually the keyword, is required to be not null and unique. The other columns are the information need to be translated. The number of columns upload in one project should be the same.

### Task division

Each entry in upload files is an independent task. The entry has to go through three stages before release: assignment, translation, review. The change of entry state is showed (Fig. 3). User needs to continuously translate until it passes the review. Only those qualified entries are released, and there is no need to wait all entries are qualified. Subsequent qualified translations can be added to the next version, and all revisions are based on the last version.

**Fig. 3.**
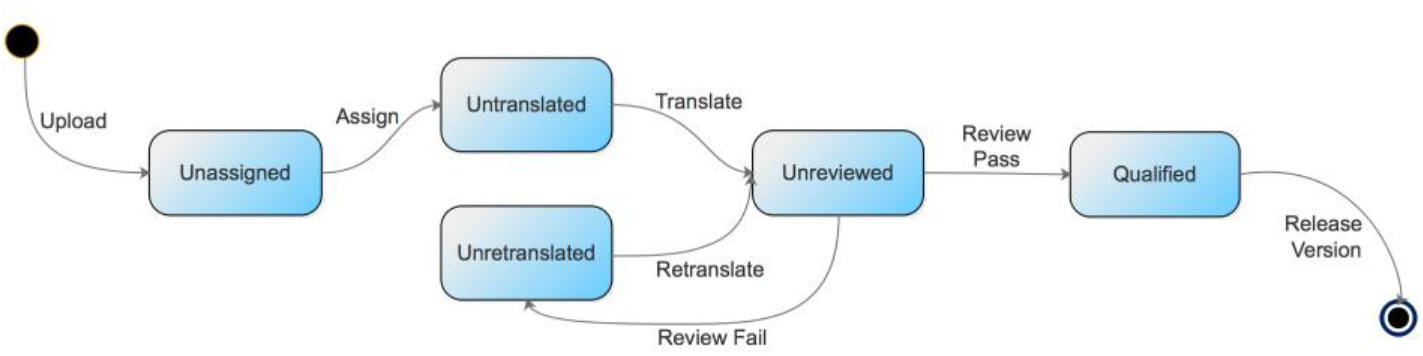
Entry State During the Translation

### Record tracing

We’ve recorded users’ every translation and related review in the database, which includes the translation, translated time, review result, and reviewed time. Furthermore, all historical uploaded files are saved on the server. All these are not easy to access and edit by general users. The system can be easily restored from disaster with these files. Regular backup of the database and uploaded folder are suggested.

### Quick Start

The programming is object-oriented and adopts the popular framework, vue and laravel, which would be easily extended and modified. The whole project is docker-packaged, thus system administrator or system operation engineer would easily integrate and deploy, isolating from complex configuration and dependency problems.

## Results and Discussion

### Build Translate Projects with Different Permissions

Each registered user could create a translation project and assign different permissions to project members. Members follow the procedure (Fig. 4) and coordinately promote the project development. Initially, the owner creates a project, uploads database file, and assigns database entries to a translator. The translator continuously translates these assigned entries with the help of Google Translate until approved. Then the reviewer checks these translations carefully one by one and decides whether to let it pass or not. Finally, the owner releases a version of qualified translations that passed the previous review. An issue is opened if found error in released translations, and the owner needs to decide whether to re-translate the entry or not. If re-translation is determined, the entry is reassigned and the previous steps are repeated. Until all translations are in good quality, then the project finished. As the permission is only related to projects, it is feasible for one user to join multiple projects with different roles. Furthermore, every logged-in user can browse released translations and open issues once the visibility level of a project is set to public. With more and more biomedical practitioners join the projects powered by Trantrace, an active community is established to update and maintain those translated databases.

**Fig. 4.**
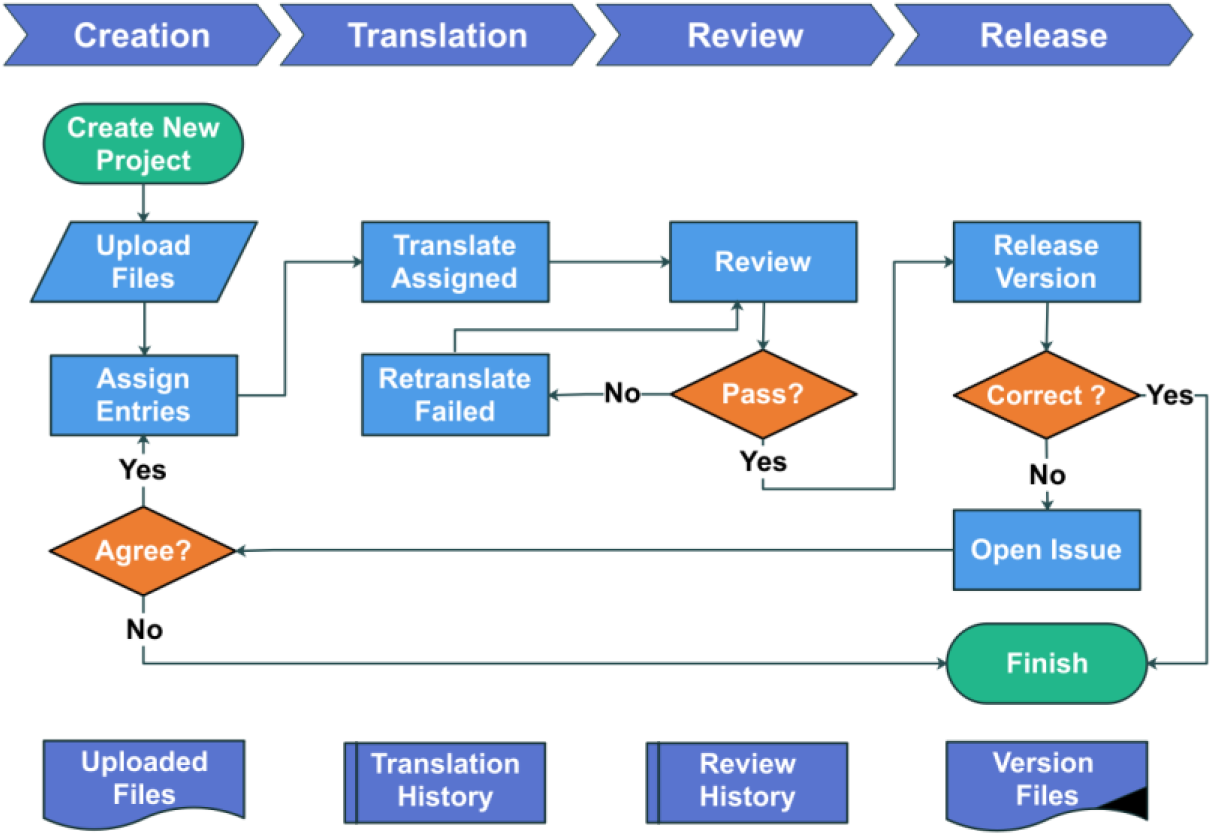
Translation Workflow in Trantrace

### Version Control Your Translation

Version control depicts the translation revolution in long-running projects. A version is released when the project owner determines that current progress is remarkable. Only qualified translations are integrated, others are excluded from this version. The follow-up optimization and correction are based on the latest version. All historical versions are recorded and downloadable. Furthermore, all translations and related reviews are recorded in the history panel, even if it is rejected by the reviewer or excluded from released versions. All these together would make Trantrace more robust for long-term collaborative translations.

### Up-to-date Task Management

Task management encompasses two parts (Fig. 5): project and personal task management, which could serve as a foundation of an efficient workflow. The up-to-date project progress is obvious through the bar chart in the project and member schedule, from which the owner can know the workload and completed the percentage of each translation stage and project member. Personal unfinished tasks are summarized in the dashboard by project and in my task by the category, respectively. The dashboard shows the deadline, priority, total task counts, completed percentage and unfinished counts of joined projects. Users could arrange their work organized and prioritized. My task broadly calculates the unfinished numbers no matter which projects it is. Those unfinished counts contain a shortcut to the corresponding workspace. Furthermore, a pop-up dialog would inform users of those tasks every minute. All these are lumped together into the basic activities and would make task management efficient.

**Fig. 5.**
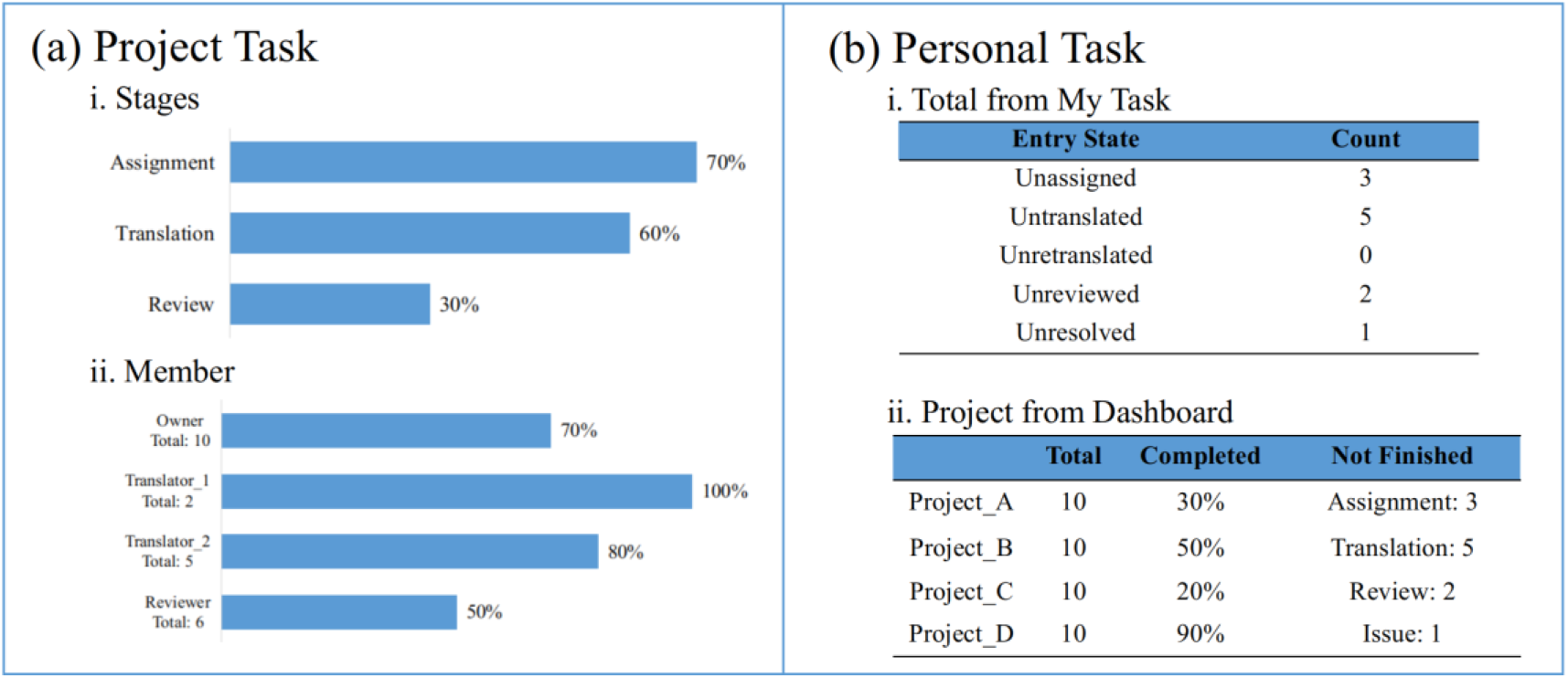
Task Management

### Limitation and future directions

Trantrace is aimed to facilitate collaborative translation rather than substitute human translation. We only provided an external link to Google Translate with copied content, since the popular translation API, such as Google Translation API (https://cloud.google.com/translate/) and Microsoft Translator Text API (https://www.microsoft.com/en-us/translator/business/translator-api/), are always charged. Given that those online biomedical resources are always large and frequently updated, it remains to be tedious for manual download. Automatic download, comparison, and upload of those well-known and public resources would be the subject of further development.

## Conclusions

Precision medicine has speeded diagnosis, influenced preventive medication and improved life quality for individual patients. To maxmize the benefit population, we proposed an open-source web-based Trantrace for continuous collaborative biomedical translation. It offers a simple way for translation cooperation and grants each user more freedom to build own team. Every translation is traceable and reproducible owing to the version control mechanism. Users could get rid of inconsistent translation content and focus on own task fulfillment. Real-time prompts would remind users of unfinished tasks to balance their work. All these together would streamline resources translation and make it more productive.

## Availability and requirements

Project name: Trantrace

Project home page: https://github.com/sgi-drylab/trantrace

Operating system: Linux, OSX

Programming language: PHP, MySQL, and Javascript

Other requirements: PHP≥7.1.3; MySQL≥5.7

License: MIT

Any restriction to use for non-academic: None

## Abbreviations

TCGA: The Cancer Genome Altas
HMP: Human Microbiome Project
ACMG: American College of Medical Genetics
AMP: Association of Molecular Pathologists
CPIC: Clinical Pharmacogenetics Implementation Consortium
TMS: Translation Management System
GUI: Graphic User Interface

## Declarations

### Availability for data and materials

Web demo of Trantrace can be accessed via http://trantrace.sgi-drylab.org. Online documentation is available at http://trantrace.sgi-drylab.org/help.

### Author’s contributions

C.W. and M.S. conceived and designed the project; Z.W., M.L., and P.C. performed the software development and initial testing under the supervision of C.W.; Z.H. and contributed to project testing and further refinement; M.S. wrote the manuscript and documentation with support from C.W.; M.H. helped with drawing of figures; J.C. provided server setting and domain hosting; C.W. and R.L. provided the resources support. All authors approved the final version of the manuscript.

## Acknowledgements

This work was supported by Singlera Genomics.

## Authors’ information

^1^Singlera Genomics Inc., Shanghai, China

